# Interpretable Pairwise Distillations for Generative Protein Sequence Models

**DOI:** 10.1101/2021.10.14.464358

**Authors:** Christoph Feinauer, Barthelemy Meynard-Piganeau, Carlo Lucibello

**Affiliations:** Department of Decision Sciences, Bocconi Institute for Data Science and Analytics (BIDSA) Bocconi University, Milan, Italy; Laboratory of Computational and Quantitative Biology (LCQB) UMR 7238 CNRS - Sorbonne Université, Paris, France; Department of Applied Science and Technologies (DISAT), Politecnico di Torino, Torino, Italy

## Abstract

Many different types of generative models for protein sequences have been proposed in literature. Their uses include the prediction of mutational effects, protein design and the prediction of structural properties. Neural network (NN) architectures have shown great performances, commonly attributed to the capacity to extract non-trivial higher-order interactions from the data. In this work, we analyze three different NN models and assess how close they are to simple pairwise distributions, which have been used in the past for similar problems. We present an approach for extracting pairwise models from more complex ones using an energy-based modeling framework. We show that for the tested models the extracted pairwise models can replicate the energies of the original models and are also close in performance in tasks like mutational effect prediction.

## 1 Introduction

Many different types of generative models for protein sequences have been explored, from pairwise models inspired by statistical physics [1, 2, 3, 4] to more complex architectures based on neural networks like variational autoencoders [5, 6, 7], generative adversarial networks [8], autoregressive architectures [9, 10] and models based on self-attention [11]. While such models promise a rich field of applications in biology and medicine [12], the question of what information they extract from the sequence data has received less attention. This is, however, a very interesting field of research since especially the more complex models might extract non-trivial higher-order dependencies between residues. This in turn might reveal interesting biological insights.

Some recent works address this interpretability issue. In Ref. [13], the authors introduce the notion of *pairwise saliency* and use it to quantify the degree to which more complex models learn structural information and how this relates to the performance in the prediction of mutational effects. Ref. [14] instead constructs pairwise approximations to categorical classifiers and showcases applications to models trained on protein sequence data.

We observe that the performance of many different models on tasks like the prediction of mutational effects is often similar even when using very different architectures and, in addition, is close to what simple, pairwise models achieve (see e.g. [9]). It appears natural to ask then how much of the predictive performance of the more complex models like variational autoencoders is due to higher-order interactions which are inaccessible to more simple models.

We therefore ask in this work how close trained neural network (NN) based models are to the manifold of pairwise distributions. To this end, we train three different architectures on protein sequence data. Interpreting these models as energy-based models [15], we present a simple way to extract pairwise models from them and analyze errors in energy between extracted and original models. We show that the subtle question of gauge invariance is important for this purpose and address this invariance ambiguity using different objective functions for the extraction.

## 2 Methods

### 2.1 Protein Sequences and Energy-Based Models

We represent the aligned primary structure of a protein domain of length *N* as a sequence *s* = (*s*_1_, …, *s*_*N*_), where we identify every possible amino acid with a number between 1 and *q*, where *q* is the number of possible symbols (we use 20 amino acids and 1 gap symbol, so *q* = 21). Our input data are sets of evolutionary related sequences gathered in multiple sequence alignments (MSAs), where every row corresponds to a sequence of amino acids and every column to a consensus position [16].

Energy-based models (EBMs) [15] are models that specify the negative unnormalized log-probability *E*_*θ*_(*s*), for example by a neural network with weights and biases represented by *θ*. While the calculation of the exact probability

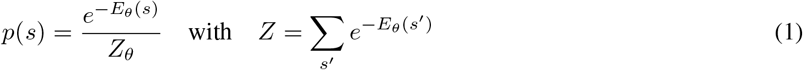

is intractable since the normalization constant *Z*_*θ*_ is a sum over *q*^*N*^ terms, numerous ways of training such models have been developed.

In this work, we use the fact that *any* probability *p*(*s*) can be thought of as an EBM by defining *E*(*s*) = −log *p*(*s*). We will use the term *energy* for both cases: when derived from a distribution *p*(*s*), which is typically normalized, and when given by an explicit energy function, which is typically not normalized. While this formulation could be extended to models for sequences of varying length, we restrict ourselves in this work to sequences of fixed length.

### 2.2 Energy Expansions and Gauge Freedom

We call *I* = {1, …, *N*} the set of all positions in the sequence *s* and *s*_*L*_ the subsequence consisting of amino acids at positions in *L* ⊆ *I*. Then, we can expand any energy *E*(*s*) in the form

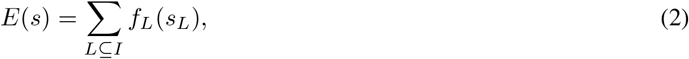

where *f*_*L*_ is a function depending only on the amino acids at positions at *L*. We will use *f* for denoting the set of all *f*_*L*_ in the expansion. Models for which *f*_*L*_ = 0 for |*L*| > 2 are called *pairwise models* (or *Potts models*) and their energy can be written as a special case of Eq. 2 as

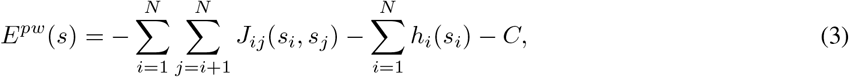

with *J* being commonly called couplings and *h* the fields [17]. The constant *C* is typically not added to the model definition since it does not change the corresponding probabilities, but we keep it in order to be consistent with the generic expansion in Eq. 2. While models for which *f*_*L*_ = 0 for |*L* | > 1 are often considered separately as *independent site* or *profile* models [9], we regard them in this work as special cases of pairwise models.

The expansion in Eq. 2 is not unique, which means that given an energy *E*(*s*) it is possible to find different expansion parameters *f* for which Eq. 2 holds. Therefore additional constraints must be imposed to fix the expansion coefficients (gauge fixing). It is for example trivial to rewrite the *pairwise* model in Eq. 3 as a model with interactions only of order *N* by defining *f*_*I*_ (*s*) = *E*(*s*) and *f*_*L*_ = 0 for |*L* |< *N*. A common route is to impose the so-called *zero-sum* gauge [14], which aims to shift as much of the coefficient mass to lower orders as possible (see, e.g., Ref[14] and Appendix B.2 for details). This is intuitively sensible, since explaining as much of the variance as possible with low order coefficients seems to be a key element when trying to understand how complex the model is. However, we will show in the next section that the problem of gauge invariance is more subtle and important for understanding the structure of the fitness landscape induced by NN models.

### 2.3 MSE Formulation

We formulate the problem of extracting a pairwise model from more general models by using a loss function ℒ that measures the average mean squared error (MSE) in energies with respect to a distribution *D* over sequences. We call *E*^*M*^ (*s*) the energy of the original model that we want to project on the pairwise space. We define the loss function over the parameters *J, h* and *C* on which the pairwise energy *E*^*pw*^(*s*) of Eq. 3 implicitly depends as

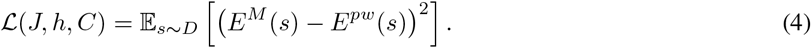

We minimize the loss function with respect to *J, h* and *C* and use the resulting pairwise model *E*^*pw*^ as an approximation to *E*^*M*^ .

The distribution *D* is central in this formulation of the problem and is closely related to the question of gauge invariance. It can be shown that if *D* is the uniform distribution over sequences, the minimizer of ℒ(*J, h, C*) is equivalent to the pairwise part of *E*^*M*^ in the zero-sum gauge (see Appendix B for a proof). This means in reverse, that extracting the pairwise model using the zero-sum gauge is equivalent to minimizing the MSE in energy when giving all possible sequences equal weight. However, generative models trained on protein families are used only on a small region of the sequence space. By changing *D* it is possible to give more weight to these regions and construct a pairwise model that might be worse in replicating *E*^*M*^ globally, but better in regions of interest. This is equivalent to extracting the pairwise interactions in a different gauge of *E*^*M*^ .

Note that if the original model is in fact a pairwise model, then for any *D* with a sufficiently large support the minimizer of Eq. 4 should correspond to the original model (up to a gauge transformation).

A natural candidate for *D* is the distribution induced by *E*^*M*^, leading to pairwise models that aim to reproduce the original distribution well on typical sequences of that distribution. With this choice, the loss corresponds to an *f*-divergence (*f* (*t*) = *log*^2^(*t*)) in the unnormalized distribution space [18]. Notice also that for a trained model *E*^*M*^, one would expect this distribution to be close to the training distribution.

In the following we test how well extracted pairwise models reproduce the energies of the original models when using different distributions *D* in Eq. 4.

## 3 Results

### 3.1 Extraction of Fourier Coefficients

We train three different probabilistic models (the autoregressive architecture presented in [9] (ArDCA), an energy based model expressed by a multi-layer perceptron with a single hidden layer (MLP) and a variational autoencoder [5] (VAE), on five different MSAs taken from [3]. The datasets correspond to mutants and experimental fitness values for the *BRCA1* tumor suppressor gene [19], the *GAL4* transcription factor [20], the poly(A)-binding protein *PABP* [21], the ubiquitination factor *UBE4B* [22] and the yes-associated protein *YAP1* [23].

After training the models we prepare 10^7^ samples from the uniform distribution U and 10^7^ samples from the model distribution M. Using the corresponding model energies we extract a pairwise model by minimizing the loss in Eq. 4 (see Appendix A.2 for details of the models and the training procedure).

### 3.2 Energy Errors

In Fig. 1 we show the error in the energies of extracted pairwise models with respect to the energies in the original models. We use two different distributions *D* in Eq. (4) for sampling the sequences used for the extraction of the pairwise models: U stands for the uniform distribution; M for the distributions of the original trained models. We evaluate the error on the sequences of the training data, test data and the mutated sequences corresponding to the experimental assays.

**Figure 1:**
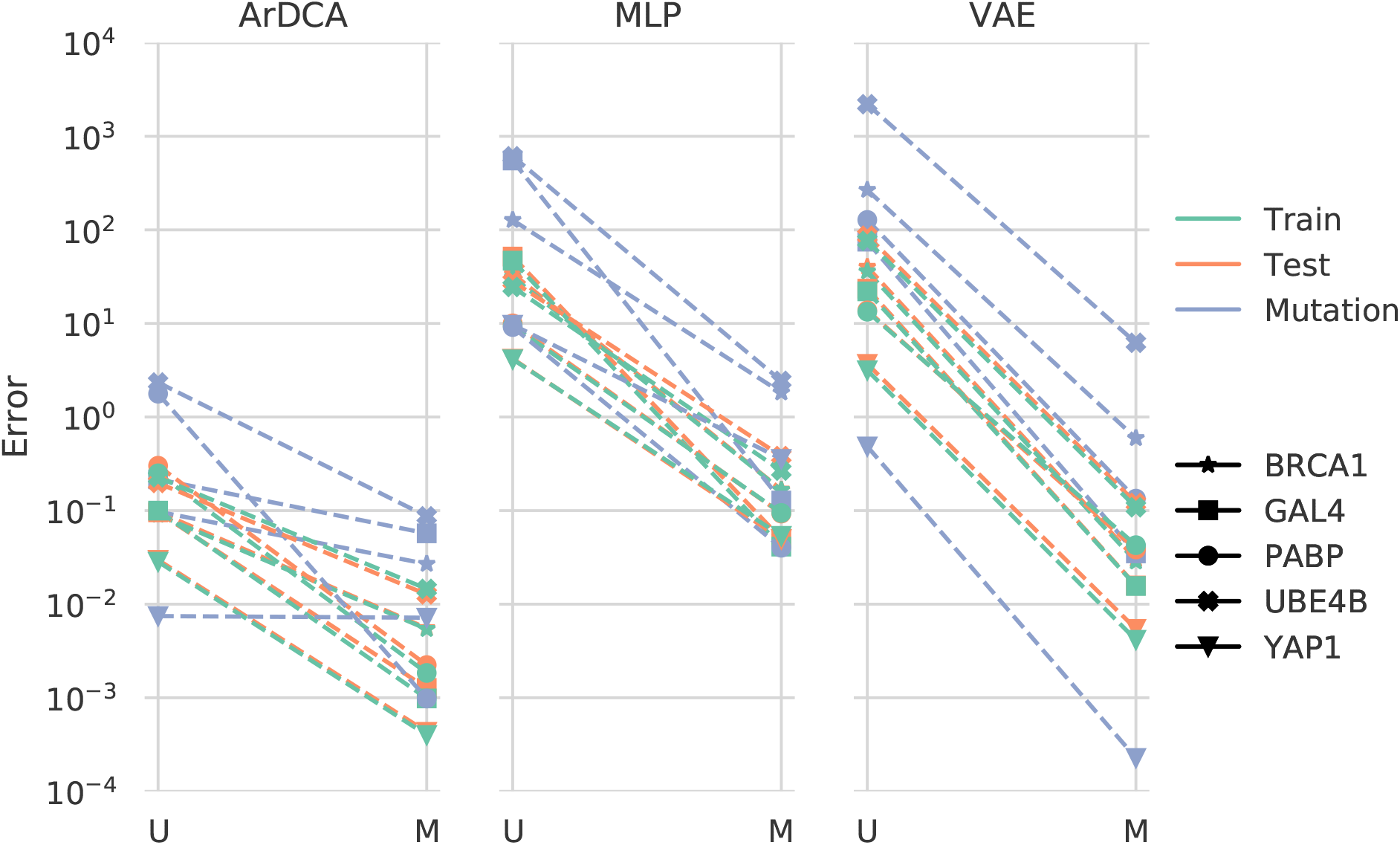
Errors in energies of the extracted pairwise models with respect to the original models. The three columns correspond to the three different models tested (ArDCA, MLP and VAE). The colors indicate which dataset is tested: Train data (green), test data (orange) and data from the mutational assays (blue). The markers distinguish the different protein families tested. Within every column, the left (U) corresponds to pairwise models extracted with samples from the uniform distribution, the right (M) to pairwise models extracted with samples from the distribution of the original models. The error shown is the normalized root-mean squared error (see Appendix A.1). Note the logarithmic scale.

The error in the plot is the mean squared error, normalized by the range (see Appendix A.1). For all models, the error drops by several orders of magnitude when using the model distribution M for extraction instead of the uniform distribution U. However, for ArDCA the error is already considerably smaller than for the other models when using the uniform distribution, which can be taken as evidence that this model is close to a pairwise distribution after training. The MLP and VAE on the other hand, show very large errors when using the uniform distribution. This can be taken as evidence that these models are either not pairwise models *globally* or that the uniform samples are not enough to extract the corresponding parameters. In both cases, the results indicate that the models are close to pairwise in the space of sequences where they are typically used.

To further highlight the difference in quality of the fit, we plot in Fig. 2 the energies of the original and extracted pairwise models after training on the PABP dataset. We evaluate the energies on the training sequences and on uniformly sampled sequences. The energy distributions between the different types of sequences are non-overlapping for all models and the energy distributions of the models on the training sequences are much wider than on uniformly sampled sequences. Furthermore, pairwise models extracted from the VAE and ArDCA with samples from the original model distribution can generalize to uniformly sampled sequences. For the MLP, on the other hand, the pairwise model extracted using sequences from the model distribution is fitting the energies on uniformly sampled sequences poorly.

**Figure 2:**
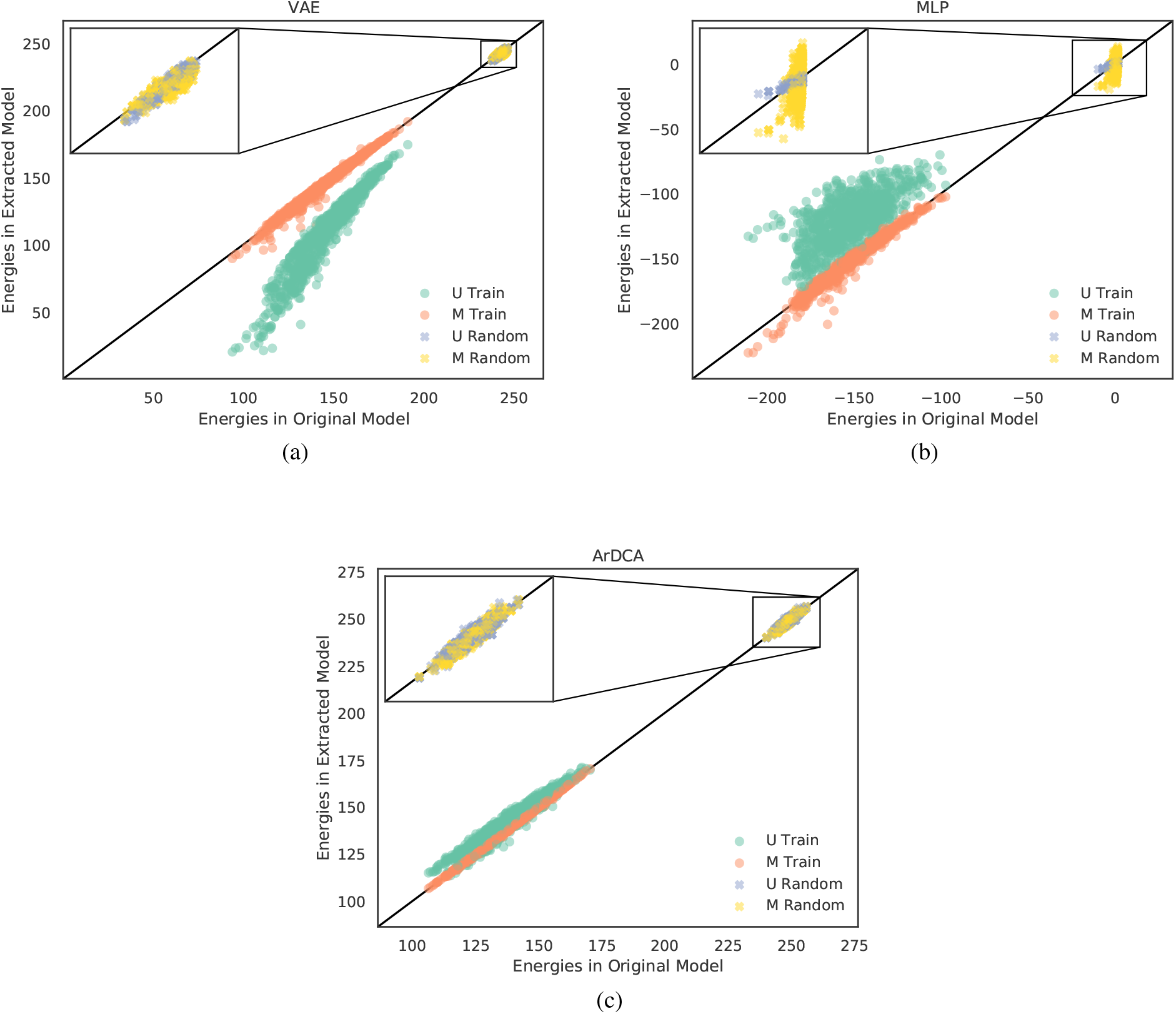
Energy Scatterplots on PABP. Plotted are energies from the original model (horizontal axis) against energies from the extracted models (vertical axis) on samples from the training data (circles) and samples from a uniform distribution (crosses). Energies are plotted for models extracted using samples from the original model distribution (red and yellow) and for models extracted using the uniform distribution (green and violet). Note that low energies correspond to high probabilities in Eq. 1. Perfect predictions would correspond to points lying on the diagonal solid line.

The energies of the training sequences are fitted well when using samples from the original model distribution for extraction. When using uniformly sampled sequences, the pairwise model extracted from the MLP overestimates the energies of the original model and the pairwise model extracted from the VAE underestimates them (note that lower energies means higher probability in Eq. 1). For ArDCA, the energies are already well fitted by pairwise models extracted with uniformly sampled sequences, with the errors further decreasing when using samples from the original model distribution. This can be taken as further evidence that the original ArDCA model, trained on this dataset, is close to a pairwise model.

### 3.3 Mutational Effect Prediction using Extracted Models

The prediction of mutational effects is a typical field of application for the type of models analyzed in this work. In Fig. 3 we show the Spearman correlations between the experimental data and the energies in the original models (O), the energies of models extracted using samples from a uniform distribution (U) and the energies extracted from the original model distribution (M). There is no clear tendency with respect to the relative performance of the original and the extracted models. This is evidence that most of the explanatory power of the original models can be reproduced by simpler pairwise models, even though the exact distribution used for extraction does not seem to be important. This is in line with the results discussed in the last section and specifically with Fig. 2: While the pairwise models extracted with uniform sequences have a significantly larger error in terms of the reproduction of the energy of training sequences, they seem to be able to largely recover the ranking of the mutated sequences in the original model.

**Figure 3:**
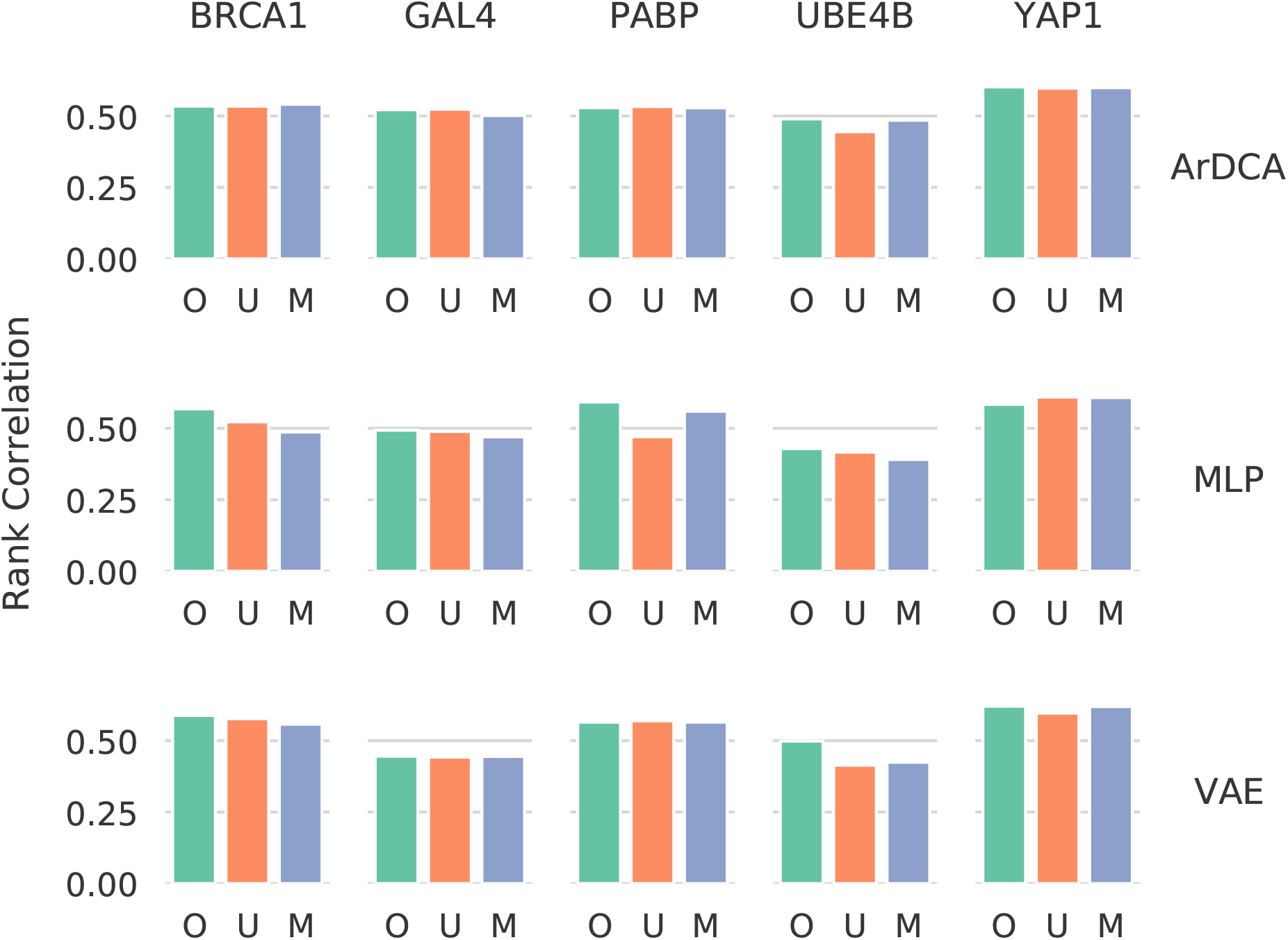
Spearman Correlation with experimental data of original (O) and extracted models (U, M). Every plot corresponds to a combination of original model type (ArDCA, MLP and VAE) with a mutational assay. Shown is the Spearman rank correlation between the experimental data and the energies of the original model (O), the model extracted using samples from a uniform distribution (U) and using samples from the original model distribution (M).

**Figure 4:**
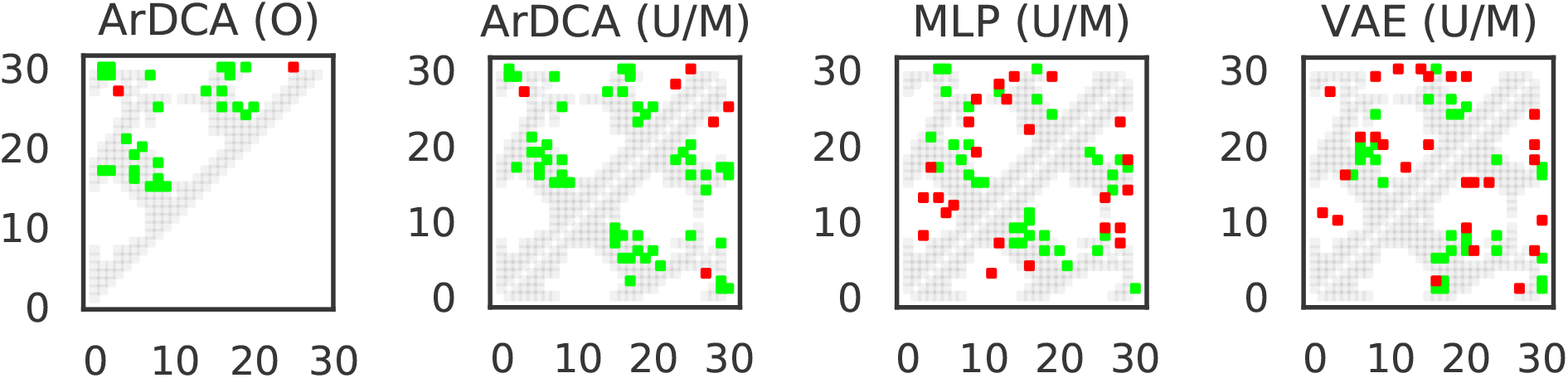
Contact prediction using extracted models. Contact predictions vs. ground truth for the top *N* = 30 predicted contacts for models extracted from ArDCA, the VAE and the MLP. Horizontal and vertical axes show positions. True contacts are grey, true positives are green, and false positives are red. In the three right plots, the upper parts show the contacts for models extracted with the uniform distribution, the lower parts show the same for models extracted with the original model distribution. The left-most plot shows the contact predictions for ArDCA from the original method in [9].

### 3.4 Contact Prediction

Given that the extracted models are pairwise models, we can use standard methods from this field to predict structural contacts [17, 24] (see Appendix A.3 for the contact prediction pipeline and the PDBs used). For ArDCA, the contact predictions for these two methods of extraction are largely the same, and also very similar to the predictions from the original method. This is consistent with the idea that ArDCA is very similar to a pairwise model. The predictions for the MLP are also very similar between the two methods and the overall performance is worse than ArDCA. The results for the VAE are similar to the MLP, indicating that the VAE and the MLP are either not relying on structural information for predicting mutational effects or our method is not able to extract this information.

## 4 Discussion

In this work, we provide evidence that the neural network based generative models for protein sequences analyzed by us can be approximated well by pairwise distributions. The autoregressive architecture on which ArDCA is based seems to be closest to a pairwise model after training. For the MLP and the VAE, the results seem to at least indicate that their pairwise projection is a very close approximation in the part of the sequence space in which they are typically used, close to the data manifold.

We cannot of course exclude that the neural network models tested by us do extract some meaningful higher-order interactions from the data, but the results seem to indicate that their effect is rather subtle. This suggests that the general strategy outlined in [25], where the pairwise part of the model is kept explicitly and an universal approximator is used for extracting higher-order interactions, might be promising. Another question is how the specific architectures and hyperparameters chosen by us influence the result. In Ref. [13], for example, the authors test many different hyperparameters for variational autoencoders, which might also have an influence on how well the resulting distributions are approximated by pairwise models.

Several interesting further lines of research suggest themselves. While the general idea of approximating a pairwise distribution over fixed-length sequences to models trained on unaligned data (like recent very large attention-based models [26]) seems to be ill-defined, the approach of extracting a pairwise model for a small part of the sequence space as highlighted in this work might still be feasible. Another interesting question is whether sparse higher-order interactions can be efficiently extracted from neural network based models. It is for example possible that methods like the Goldreich-Levin algorithm [27] might be adapted for pseudo-boolean functions based on generative models for protein sequence data.

## A Methods

### A.1 Energy Error

We measure the error in energies in the extracted models with respect to the energies in the original models using the *normalized root-mean square deviation*, i.e.

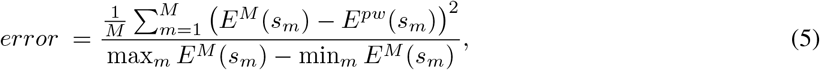

where 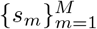 is the set of sequences on which we calculate the error, *E*^*M*^ is the energy of the original model, *E*^*pw*^ the energy of the extracted pairwise model and max_*m*_ *E*^*M*^ (*s*_*m*_) and min_*m*_ *E*^*M*^ (*s*_*m*_) are the maximum and minimum energies of the original model on the dataset.

### A.2 Models and Sampling

#### A.2.1 ArDCA

The model used in ArDCA [9] decomposes the probability *p*(*s*) of a sequence of amino acids as

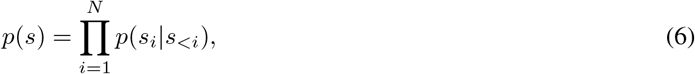

where *s*_*i*_ is the amino acid at position *i* and *s*_<*i*_ are the amino acids that come before *i* in the ordering. The conditional probability *p*(*s*_*i*_|*s*_*<i*_) is then defined as

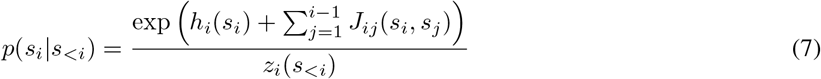

where *z*_*i*_(*s*_*<i*_) is the sum of the denominator over all possible values of *s*_*<i*_. We use the code by the authors for training the model and calculating log *p*(*s*) for the samples used for extraction. Training was done with sequence reweighting as implemented by the authors of [9].

#### A.2.2 MLP

The MLP is a simple feed-forward network with one hidden layer of size *H*. The energy *E*^*MLP*^ for sequence *s* is calculated as

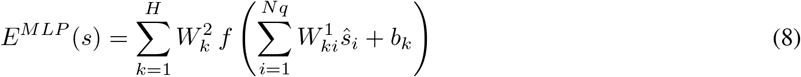

where *ŝ* is a one-hot encoding of the sequence *s, W* ^1^ and *W* ^2^ are a weight matrix and a weight vector respectively, and *b* is the bias vector. The activation function *f* was chosen as the leaky ReLU [28]. We used *H* = 64 hidden units, a L2 regularization of 0.001. The training was done using Pseudolikelihoods inspired by [24]. See [25] for the definition of the loss function when using EBMs on proteins. We use the same sequence reweighting technique as for the VAE (see next section). Training was done for 200 epochs. After training, the energy can be calculated using a single forward pass. For sampling from this model, we resorted to standard MCMC techniques [29]. Since we have to evaluate the energy a large number of times during sampling, we used a very small number of MC sweeps (MC steps divided by the length of the sequence) for thermalization (1000 sweeps) and sampling (every 5 MC sweeps after thermalization). While this certainly does not lead to high-quality samples, we note that we are only interested in biasing the extraction towards sequences more typical of the distribution. The model was implement in PyTorch [30].

#### A.2.3 Variational Autoencoder

The model and code we use is based on the work and implementation of [5]. For a more detailed introduction to the variational autoencoder we refer to the original work [31]. Both encoder and decoder use a single hidden layer with 100 hidden neurons and *tanh* activations. The dimension of the latent space is 20. During training, an *L*2 regularization of 0.1 was used and the training was run for 10000 epochs. Following the implementation of [5], we used full-batch gradient descent with an Adam optimizer.

The probabilities were estimated using importance sampling [32] using 5000 ELBO samples. Training was done with sequence reweighting as implemented by the authors of [5].

#### A.2.4 Extraction

We use 10^7^ samples from the uniform and 10^7^ samples from the original model distributions after training for extracting the pairwise models. The samples are drawn independently for each combination of original model and dataset.

For the samples from the model distributions, we minimize the loss in Eq. 4 using a batch size of 10000 and the Adam optimizer [33]. We keep a running average *l* of the loss function using the equation *l*_*k*_ = *α l*_*k* −1_ + (1 − *α*) ℒ_*k*_ with the initial condition *l*_1_ =ℒ_1_ where ℒ_*k*_ is the loss after gradient step *k* and *l*_*k*_ is the running average of the loss after gradient step *k*. We set *α* = 0.1 and stop optimization if the running average has not reached a new minimum for 1000 gradient steps. The extraction runs in seconds to minutes on an Nvidia RTX 2080 GPU.

For samples from the uniform distribution, the minimizer of the loss in Eq. 4 can be calculated directly from the samples without gradient descent (see Appendix B). We use the same number of samples to approximate the conditional energy expressions in Eq. 14 this case. We found that in the samples from the model distributions *M* not all amino acids were present in all positions. We therefore add 1% samples from the uniform distribution to the samples from the model distributions.

### A.3 Contact Prediction

We use standard methods for contact prediction from pairwise models, following mainly [24]. We transform the extracted pairwise models into the zero-sum gauge and calculate the Frobenius norm of the *q* − 1 × *q* − 1 submatrices *J*_*ij*_ corresponding to the pair of positions *i* and *j* (we do not sum over gap states, hence *q* − 1 instead of *q*). We apply the *average-product correction* [34] and sort the positions pairs by the resulting score, excluding pairs for which *abs*(*i* − *j*) *<* 5. We map PDB 1PIN:A [35] to the MSA and use it to differentiate contacts from non-contacts (8 Å, Heavy-Atom criterion [17]).

## B Zero-Sum Gauge

In the following we prove that the pairwise model *E*^*pw*^ corresponding to the minimizer of Eq. 4 is equivalent to the pairwise part of *E*^*M*^ in the zero-sum gauge when using the uniform distribution D for extraction.

### B.1 Notation

We denote by 𝒜={1, ..,*q*} the (numeric) alphabet of the *q* possible amino acids. The terms *f*_*L*_ : 𝒜 ^|*L*|^ → ℛ in the general expansion in Eq. 2 are functions mapping sequences of amino acids of length |*L*| to a real number, where *L* ⊆ *I* = {1, …, *N*} is a subsequence of positions. In this notation, the pairwise model we train using the loss in Eq. 4 can be written as

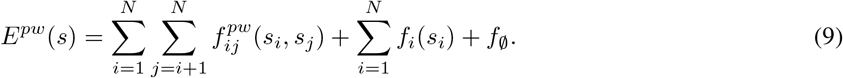

In Eq. 3 we use a different notation for the pairwise model, but in this Appendix we decide to keep all notations compatible with the generic expansion in Eq. 2. The notations can be connected by identifying 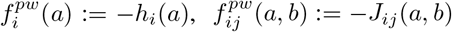 and *f*_∅_ := *−C* for arbitrary amino acids *a* and *b*.

Equivalently we define 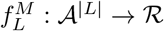 as the interaction coefficients between the sites belonging to the set of positions *L* ⊆ *I* in *E*^*M*^ in a certain gauge.

We will use *f*_*L*_(*a*_*L*_) in order to denote a specific interaction coefficient for a fixed sequence of amino acids *a*_*L*_ of length |*L*|, for both pairwise models and models with higher-order interactions. We will use *f*^*pw*^ to denote the set of all parameters of the pairwise model and *f* ^*M*^ for the set of all parameters of the original model.

### B.2 Zero-Sum Gauge

The zero-sum gauge is a reparameterization of the interaction coefficients which leaves the energy invariant (see also Ref. [14] who discuss this gauge). In this gauge, if |*L*| > 0, summing *f*_*L*_(*a*_*L*_) over any of the amino acids in *a*_*L*_ while keeping the others fixed is 0. It can be applied both to the parameters of the extracted pairwise model *f*^*pw*^ and the parameters *f* ^*M*^ of the original model. Since the sum over an amino acid is proportional to the expectation of *f*_*L*_(*a*_*L*_) when the corresponding amino acid is sampled uniformly, this condition can be written as

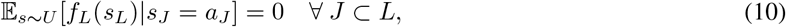

where 𝔼_*s∼U*_ [*f*_*L*_(*s*_*L*_)|*s*_*J*_ = *a*_*J*_] the expectation of *f*_*L*_(*s*_*L*_) if the subsequence *s*_*J*_ is fixed to *a*_*J*_. Any model can be transformed into the zero-sum gauge using the identity 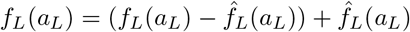 with

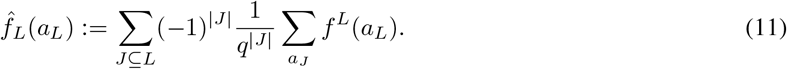

It is easy to show that 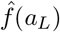 satisfies the condition in Eq. 10 and that 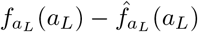 contains only interactions of order strictly less than |*L*|. Therefore, any model can be transformed into the zero-sum gauge by first applying the transformation to the interaction coefficients at the highest order *N* = |*I* |. This will lead to interaction coefficients at order *N* that satisfy the condition in Eq. 10 and new interaction coefficients of order lower than *N*. These can be absorbed in the interaction coefficients in the lower orders of the expansion. Repeating this procedure at *N −* 1, then at *N −* 2 etc. leads to a final model where all interaction coefficients of all orders satisfy the condition in Eq. 10.

Since the expansion of *E*^*M*^ has exponentially many interaction coefficients in general, this procedure has no practical use in our setting. However, in the next section we show that the lower orders of *E*^*M*^ in the zero-sum gauge representation can be extracted with a simple sampling estimator.

### B.3 Proof of Equivalence of Minimizer of Loss and Zero-Sum Gauge

The partial derivative of the loss in Eq. 4 with respect to a parameter 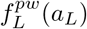 in the pairwise model (note that |*L*| ≤ 2 in this case) can be written as

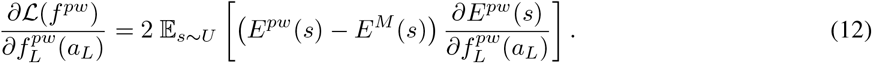

Setting the gradient to 0 leads to

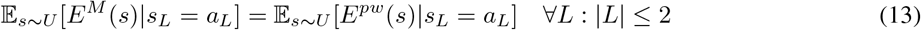

which means that the minimisation of the loss with respect to the parameters of the pairwise model is equivalent to fitting the conditional expectation of the energy under uniform distribution up to the second order of the expansion.

Since the loss in Eq. 4 is invariant with respect to a gauge change in the pairwise model *E*^*pw*^, we can assume without loss of generality that we extract the pairwise model in the zero-sum gauge representation. Using a hat to denote the parameters 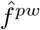 of the pairwise model in this specific gauge, it is easy to see from Eq. 9 and the condition in Eq. 10 that

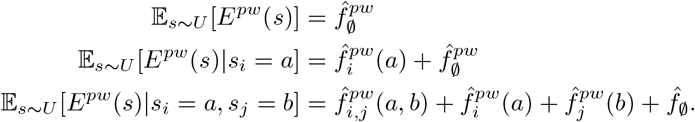

Combining this with Eq. 13 we get at the minimum of the loss the conditions

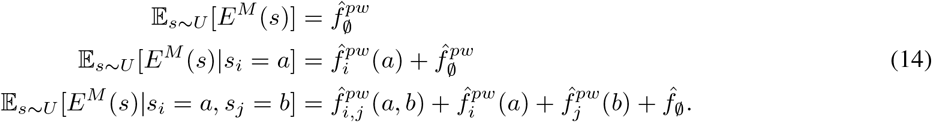

Similar to the pairwise model, we will use a hat to denote the parameters 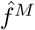 of the model *E*^*M*^ in the zero-sum gauge. While the corresponding expansion

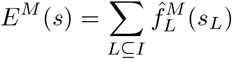

has interaction coefficients of all orders, we can again use the conditions in Eq. 10 to arrive at

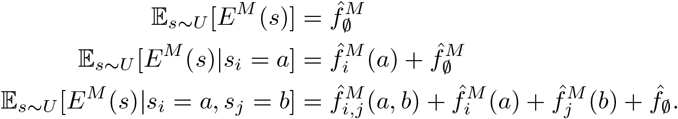

Taking these relations together leads to the minimizer condition

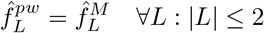

which means that the *E*^*pw*^ minimizing the loss in Eq. 4 is the pairwise part of *E*^*M*^ in its zero-sum gauge representation. Note that the loss is still invariant with respect to a gauge change in the extracted pairwise model, so the extracted model can be in any gauge representation.

We also note that Eqs. 14 can be used to estimate the coefficients of the extracted pairwise model directly using uniform samples and the corresponding energies from the original models in order to approximate the expectations.

